# Coupled myovascular expansion directs cardiac growth and regeneration

**DOI:** 10.1101/2021.01.20.425322

**Authors:** Paige DeBenedittis, Anish Karpurapu, Albert Henry, Michael C. Thomas, Timothy J. McCord, Kyla Brezitski, Anil Prasad, Yoshihiko Kobayashi, Svati H. Shah, Christopher D. Kontos, Purushothama Rao Tata, R. Thomas Lumbers, Ravi Karra

## Abstract

**Background:** Heart regeneration requires multiple cell types to enable cardiomyocyte (CM) proliferation. How these cells interact to create growth niches is unclear. Here we profile proliferation kinetics of cardiac endothelial cells (CECs) and CMs in the neonatal mouse heart and find that they are spatiotemporally coupled.

**Methods and Results:** We show that coupled myovascular expansion during cardiac growth or regeneration is dependent upon VEGF-VEGFR2 signaling, as genetic deletion of *Vegfr2* from CECs or inhibition of VEGFA abrogates both CEC and CM proliferation. Repair of cryoinjury displays poor spatial coupling of CEC and CM proliferation. Boosting CEC density after cryoinjury with virus encoding *Vegfa* enhances regeneration. Using Mendelian randomization, we demonstrate that circulating VEGFA levels are positively linked with human myocardial mass, suggesting that *Vegfa* can stimulate human cardiac growth.

**Conclusions:** Our work demonstrates the importance of coupled CEC and CM expansion and reveals a myovascular niche that may be therapeutically targeted for heart regeneration.

## INTRODUCTION

In many organs, heterologous cell types establish specialized microenvironments, or niches, that mediate tissue growth and regeneration. Alterations to niche constituents can affect the efficiency of tissue growth and outcomes after injury, making niches a possible target for therapeutic regeneration (1, 2). For organs like the intestines, bone marrow, skin, and skeletal muscle, niches are classically centered around a stem cell compartment (3). With a few notable exceptions, niches within organs that lack a resident stem cell are less defined. For instance, in the developing and regenerating neonatal mouse heart, hypoxic niches have been associated with regionalized growth, but the cellular makeup of these niches is unclear (4).

Studies in zebrafish, salamanders, and neonatal mice have established a template for innate heart regeneration through proliferation of spared cardiomyocytes (CMs) (5–8). However, innate heart regeneration is a multicellular process with required contributions from epicardial cells, inflammatory cells, and nerves (9–11). Recent work has highlighted a critical role for the vasculature in heart regeneration (12–15). In zebrafish and neonatal mice, cardiac endothelial cells (CECs) rapidly respond to injury, extending nascent vessels into the wound that ultimately guide CM growth (14, 16, 17). Lineage tracing studies have demonstrated that these new vessels form by proliferation of CECs (16–18). Functional interference with angiogenic responses in the zebrafish or the neonatal mouse heart is associated with defects in CM proliferation (14, 17). Conversely, overexpression of the master angiogenic factor, *vegfaa*, is sufficient to induce ectopic cardiac growth in zebrafish, suggesting that a stimulated vasculature instructs cardiac growth (19). However, a similar role for VEGFA-stimulated CECs in the mammalian heart has yet to be shown.

Here, we spatiotemporally model CM and CEC proliferation in the neonatal mouse heart to investigate the mechanisms underlying cardiac growth and regeneration. We find that CM and CEC proliferation are tightly coupled during postnatal cardiac growth. With cryoinjury (CI), a model of incomplete regeneration, this coupling is spatially impaired. We demonstrate that coupled myovascular expansion is dependent on VEGFA signaling to endothelial VEGFR2 and restoration of coupling after CI through exogenous *Vegfa* can enhance regeneration. Similarly, we find that genetically-determined levels of circulating VEGFA are associated with higher myocardial mass in humans, suggesting that VEGFA also regulates human myocardial growth. Together, these data demonstrate that coupled expansion of CECs and CMs within a myovascular niche regulates cardiac growth and regeneration.

## RESULTS

### Spatiotemporal coupling of EdU^+^ CECs and EdU^+^ CMs during postnatal growth

Neonatal mice are able to regenerate their hearts after injury during the first few days of life (20). The loss of regenerative capacity in the mouse heart is coincident with a developmental decline in the rate of CM proliferation. While the kinetics of CM proliferation in neonatal mice have been well-documented, dynamics of other cardiac cell types are not well characterized. To better define the relationship of CMs and CECs during the regenerative window of the neonatal mouse heart, we assayed CM and CEC cycling kinetics at various time points following an EdU pulse (Figure 1a-b). Cardiac sections at different developmental time points were stained for EdU incorporation along with PCM1 and Erg to specifically mark CM and CEC nuclei, respectively (17, 21–23). We developed customized image segmentation routines to objectively quantify large numbers of CECs and CMs from cardiac sections (3120 ± 734 CECs per heart and 2328 ± 751 CMs per heart, mean ± SD). Consistent with prior reports, we found the percentage of EdU^+^ CMs to sharply decline over the first 10 days of life, with a second peak occurring at P5 (Fig. 1b,d). The increase in EdU^+^ CMs at P5 likely coincides with a terminal round of DNA synthesis and binucleation of CMs (23, 24). EdU incorporation by CECs also declines from P1 to P10 (Fig. 1a,c) with a second peak at P7. While CM and CEC kinetics differ with regards to the timing of this second peak, the overall trends of their kinetics parallel each other. To determine the strength of this relationship, we compared rates of EdU^+^ CECs and CMs for individual hearts and found them to be correlated (R = 0.59, p = 0.0007) (Fig. 1e). Above a threshold of ∼8% EdU^+^ CECs, each 1% increase in EdU^+^ CMs is associated with a 1.03 ± 0.27% (p = 0.0007) increase in EdU^+^ CECs.

**Figure 1.**
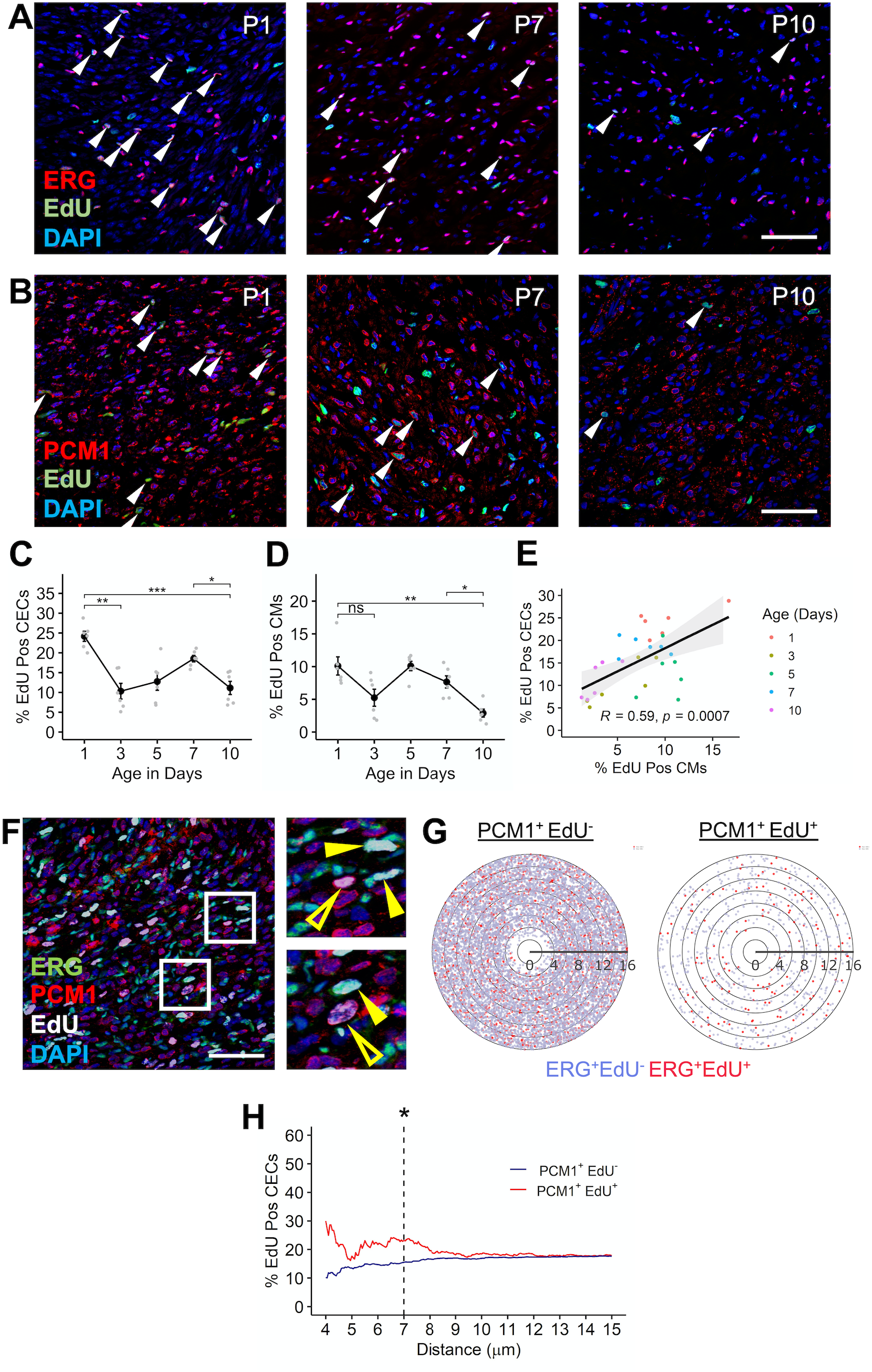
Myovascular coupling during early neonatal growth. **(A)** Representative images from postnatal day 1 (P1), 7 (P7), and 10 (P10) neonatal mouse hearts immunostained for Erg to mark CECs and EdU to identify proliferating CECs (arrows). **(B)** Representative images immunostained for PCM1 and EdU. Arrows point to EdU^+^PCM1^+^ proliferating CMs. **(C, D)** Quantitation of CEC and CM proliferation (n = 6 mice per time point). Each gray point is an individual heart. *p < 0.05, **p < 0.01, ***p< 0.001, ns = not significant, two-sided t-test corrected for multiple comparisons using Holm’s method. **(E)** Correlation of CEC and CM proliferation. Black line is the best-fit regression line and gray area indicates the 95% CI. p value indicates significance of Pearson correlation. Each point represents an individual heart and is color coded by age. **(F)** Representative image from a P4 heart immunostained for PCM1, Erg, EdU, and DAPI. Boxed regions correspond to the adjacent magnified panels. Hollowed, yellow arrows are PCM1^+^EdU^+^ CMs and solid yellow arrows are Erg^+^EdU^+^ CECs. **(G)** Pseudodistance maps for detecting enrichment of EdU^+^ CECs around EdU^+^ CMs. Distance information for 12,374 CEC-CM pairs from 6 mice was used to map EdU^+^ CECs (red dots) and EdU^-^ CECs (blue dots) relative to an averaged EdU^-^ and EdU^+^ CM (center of the plot). Each line represents 2 μm from the center of an averaged CM nucleus. **(H)** Proportion of CEC’s that are EdU^+^ as a function of distance (μm) from PCM1^+^EdU^+^ nuclei (red) and PCM1^+^EdU^-^ nuclei (blue). 274 EdU^+^ CMs and 3658 EdU^-^ CMs from 6 mice were considered, p = 0.04 at 7 μm, two-sided Z-test. Scale bars are 50 μm.

Based on the strong temporal association of CEC and CM proliferation, we next sought to spatially relate CEC and CM cycling. Within thick sections of hearts from P4 mice pulsed with EdU, we observed numerous instances of EdU^+^ CMs adjacent to EdU^+^ CECs (Fig. 1f). To quantify this observation, we assigned coordinates to each CEC and CM nucleus and computed pairwise distances for every CEC and CM. We used this distance information to deconvolve overlapping microenvironments by mapping the position of every CEC relative to an EdU^+^ CM or an EdU^-^ CM as a function of distance, resulting in “pseudodistance maps” (Fig. 1g). When comparing the density of EdU^+^ CECs relative to an EdU^+^ CM or EdU^-^ CM, we found the percentage of EdU^+^ CECs to be enriched around EdU^+^ CMs compared to EdU^-^ CMs until about ∼7 μm (p = 0.04 at 7 μm, two-sided Z-test) (Fig. 1g-h), providing evidence for coupled myovascular expansion during physiologic growth.

### Coupling of myovascular growth after cryoinjury to the neonatal mouse heart

Unlike other models of injury to the neonatal mouse heart, cryoinjury (CI) results in incomplete or inefficient regeneration (25–28). Detailed analyses of CM proliferation kinetics after CI have demonstrated that CM proliferation occurs in this model, but ostensibly at levels lower than that of the uninjured neonatal heart (25, 27). Thus, we chose to evaluate myovascular expansion following CI at P1 as a model of inefficient regeneration, in a regeneration-competent context. We first profiled CM and CEC kinetics, considering on average of 2407 ± 891 CECs and 2153 ± 1495 CMs within the border zone for each heart (mean ± SD). We found that CM and CEC kinetics generally follow the same trend as in the uninjured heart, with an approximately 50% decrease in proliferation indices for both cell types over the first 10 days of life (Fig. 2a-d). When we compared rates of CM and CEC proliferation in individual hearts, we once again noted that CM and CEC proliferation rates are correlated with ∼ 1% increase in the percentage of EdU^+^ CECs for every percent increase of EdU^+^ CMs (R = 0.54, p = 0.008) (Fig. 2e). However, when we performed a spatial analysis of CECs and CMs within the border zone, we did not detect enrichment of EdU^+^ CECs in the immediate vicinity of EdU^+^ CMs (Fig. 2g,h). In fact, EdU^+^ CECs may even be depleted around EdU^+^ CMs in the border zone after CI, suggesting that CEC and CM proliferation may not be efficiently coupled after CI.

**Figure 2.**
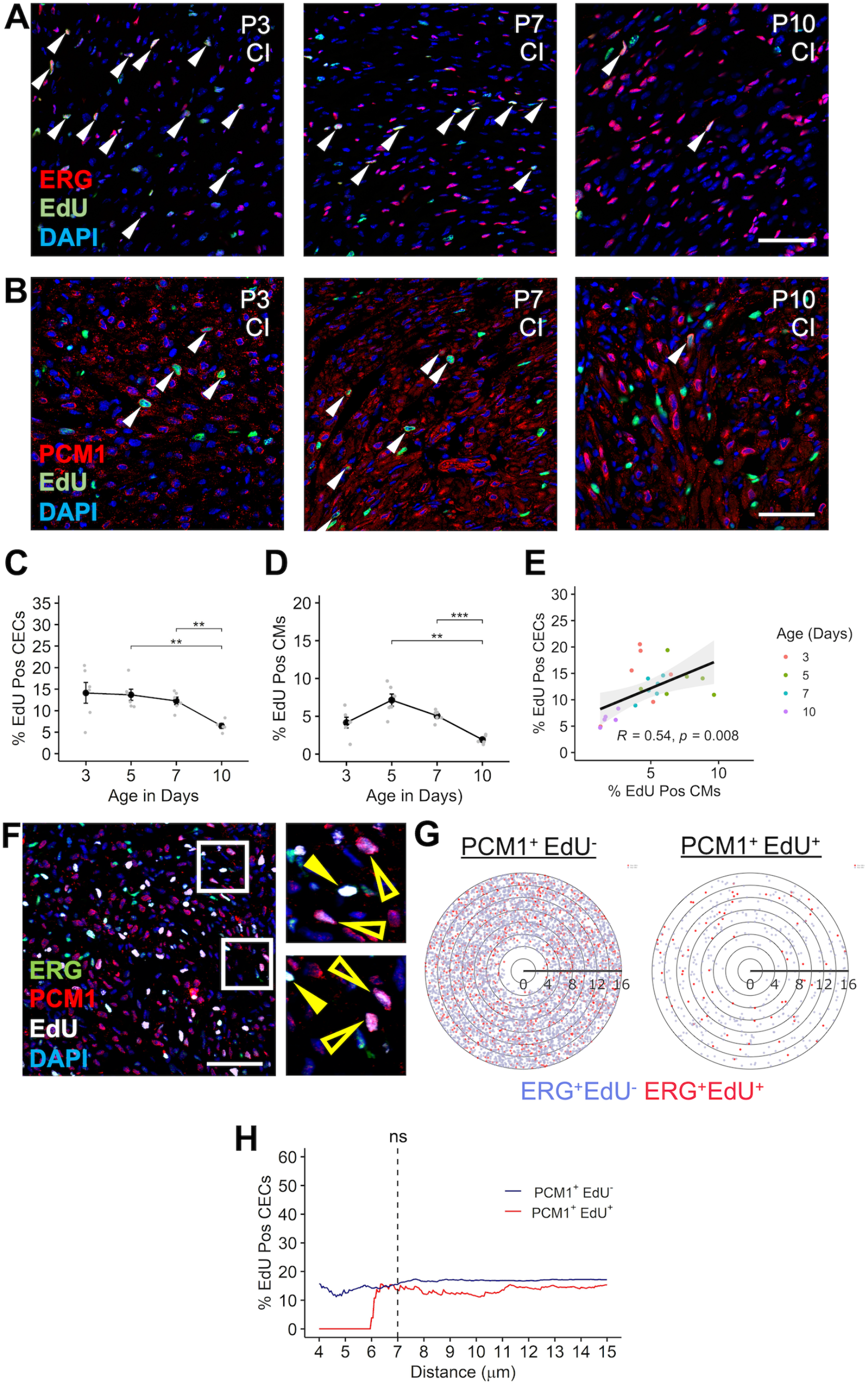
Myovascular coupling after cryoinjury. (**A, B)** Representative images from hearts collected at P1, P7, and P10 after cryoinjury (CI) at P1. Arrows point to EdU^+^Erg^+^ proliferating CECs in (A) and EdU^+^PCM1^+^ proliferating CMs in (B). **(C, D)** Quantitation of CEC and CM proliferation after CI (n = 5-6 mice per time point). Each gray point is an individual heart. **p < 0.01, ***p< 0.001, two-sided t-test corrected for multiple comparisons using Holm’s method. **(E)** Correlation of CEC and CM proliferation. Black line is the best-fit regression line and gray area indicates the 95% CI. p value indicates significance of Pearson correlation. Each point is an individual heart, color coded by age. **(F)** Representative section from the border zone of a cryoinjured P4 heart immunostained for PCM1, Erg, EdU, and DAPI. Boxed regions correspond to the adjacent magnified panels. Hollowed, yellow arrows are PCM1^+^EdU^+^ CMs and solid yellow arrows are Erg^+^EdU^+^ CECs. **(G)** Pseudodistance maps for detecting enrichment of EdU^+^ CECs around EdU^+^ CMs. Distance information for 7,093 CEC-CM pairs from 6 mice post CI was used to map EdU^+^ CECs (red dots) and EdU^-^ CECs (blue dots) relative to an averaged EdU^-^ and EdU^+^ CM (center of the plot). Each line represents 2 μm from the center of an averaged CM nucleus. **(H)** Proportion of CEC’s that are EdU^+^ as a function of distance in μm from PCM1^+^EdU^+^ nuclei (red) and PCM1^+^EdU^-^ nuclei (blue). Solid line indicates the point estimate. 186 EdU^+^ CMs and 2446 EdU^-^ CMs were considered, p = not significant (ns) at 7 μm, two-sided Z-test. Scale bars are 50 μm.

### Association of myovascular coupling with VEGFA-VEGFR2 signaling

To better understand the molecular mediators of myovascular coupling during growth and injury, we performed single cell RNA-sequencing (scRNA-Seq) using border zones of P7 hearts that were cryoinjured at P1. We profiled 1721 cells and identified 9 clusters of cells carrying markers of CECs, CMs, fibroblasts, and inflammatory cells (Supplemental Table 1, Fig. 3a). Clusters 1, 4, and 6 were notable for cells with CEC markers, such as *Fabp4, Pecam1, Erg,* and *Vegfr2* (Supplemental Table 1). Compared to the other CEC clusters, cluster 4 had increased expression of numerous proliferative markers, including *Ki67, Prc1, Ccna2, and Ccnb2* (Supplemental Table 2). A similar cluster of highly proliferative CECs has been also reported by other groups after coronary ligation in neonatal and adult mice (29, 30). In the neonatal mouse heart, VEGFR2 marks cardiac microvascular cells and these cells have recently been implicated in revascularization following injury (17). Evaluation of sections from neonatal hearts under physiologic growth conditions and after injury revealed multiple clusters of EdU^+^ CMs in close proximity to EdU^+^ VEGFR2^+^ CECs (Fig. 3b,c), supporting a role for VEGFR2^+^ microvascular CECs as contributing to a myovascular niche of growth.

**Figure 3.**
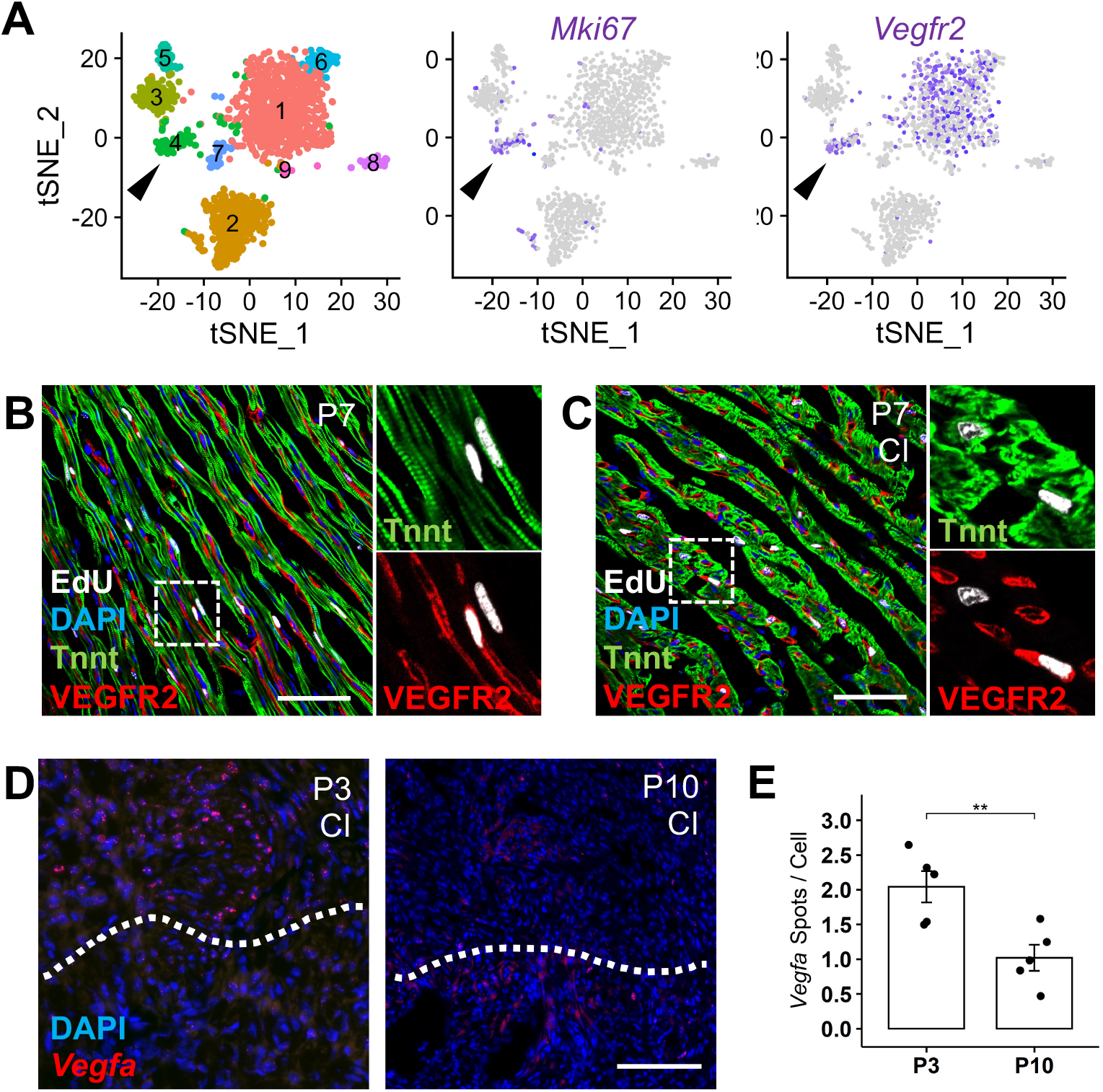
VEGFA-VEGFR2 signaling is associated with myovascular coupling. **(A)** tSNE plots of scRNA-seq data from P7 hearts that were underwent CI at P1. In the left most panel, cells are colored by clusters identified by PCA analysis. Arrows point to a cluster of CECs that are enriched for expression of *Mki67* and *Vegfr2* (purple). **(B, C)** Clusters of EdU^+^VEGFR2^+^ CECs adjacent to EdU^+^Tnnt^+^ CMs at P7 during growth (B) and after CI at P1 (C). Insets are magnifications of the regions in the solid white boxes. **(D)** Single molecule in situ hybridization for *Vegfa* in P3 and P10 hearts that underwent CI at P1. Dashed line indicates the approximate injury plane, with the border zone above the line. Red marks *Vegfa* signal. **(E)** Quantitation of in situ hybridization for *Vegfa* expression in the border zone of P3 (n = 5) and P10 (n=5) hearts after CI at P1, **p = 0.008, two-sided t-test. Scale bars are 50 μm.

For many organs, parenchymal cells secrete angiogenic factors that enable matching of vascular supply to organ size (31). Based on our prior work linking *vegfaa* overexpression to ectopic cardiomyogenesis in the zebrafish heart, we assayed *Vegfa* expression during growth and regeneration by single molecule fluorescent in situ hybridization (19). We found that under physiologic growth conditions, *Vegfa* is expressed in CMs but at low levels (Supplemental Fig. 1a). After injury, *Vegfa* is sharply upregulated throughout the ventricle (Supplemental Fig. 1b). However, based on our time course of CI responses, CEC proliferation wanes in the border zone over time (Fig. 2a). To determine whether *Vegfa* levels may be contributing to CEC dynamics after CI, we performed quantitative in situ hybridization of the border zone at P3 and P10 in hearts cryoinjured at P1 (32). We found *Vegfa* expression to decline by ∼50% from P3 to P10 in the border zone (Fig. 3 d,e), paralleling kinetics of CEC cycling. Together, these data support a dynamic role for myocardial VEGFA to endothelial VEGFR2 signaling as a regulator of the myovascular expansion during growth and regeneration.

### Requirement of endothelial *Vegfr2* for CM proliferation during growth and regeneration

Based on our scRNA-Seq experiments indicating that many proliferating CECs are *Vegfr2*^+^, we hypothesized that VEGFR2 signaling is a critical mediator of myovascular growth in the early neonatal period. To conditionally delete *Vegfr2* from CECs, we crossed *Vegfr2*^flox/flox^ mice to *Cdh5-CreER*^T2^ mice to generate *Cdh5-CreER*^T2^; *Vegfr2^flox/flox^* (*Vegfr2*^ΔEC^) and *Vegfr2^flox/flox^* (*Vegfr2^WT^*) mice (33, 34). We verified loss of VEGFR2 from CECs by immunostaining for VEGFR2 in *Vegfr2*^ΔEC^ mice (Supplementary Fig. 2a). Evaluation of CECs in *Vegfr2*^ΔEC^ mice revealed fewer CECs than in *Vegfr2^WT^* animals and a nearly 80% reduction in CEC proliferation (Fig. 4a,b). To determine how CM growth is affected by the absence of *Vegfr2* from CECs, we next assayed EdU incorporation by CMs. While CM numbers were largely preserved, CM cycling was also attenuated by ∼ 80% in *Vegfr2*^ΔEC^ mice compared to *Vegfr2^WT^* mice (Fig. 4c,d) . We next evaluated the effect of *Vegfr2* deletion from CECs on myovascular growth after cryoinjury. Similar to our results during early neonatal growth, relative numbers of CECs were decreased while relative CM numbers were preserved in *Vegfr2*^ΔEC^ hearts. We identified defects in both CEC and CM incorporation of EdU after injury in *Vegfr2*^ΔEC^ mice (Fig. 4e-h). Specifically, we observed *Vegfr2*^ΔEC^ hearts to have an almost 80% decrease in cycling CECs after injury and a 40% decrease in cycling CMs. Consistent with our hypothesis, CM expansion during growth and regeneration is dependent on intact *Vegfr2* in CECs.

**Figure 4:**
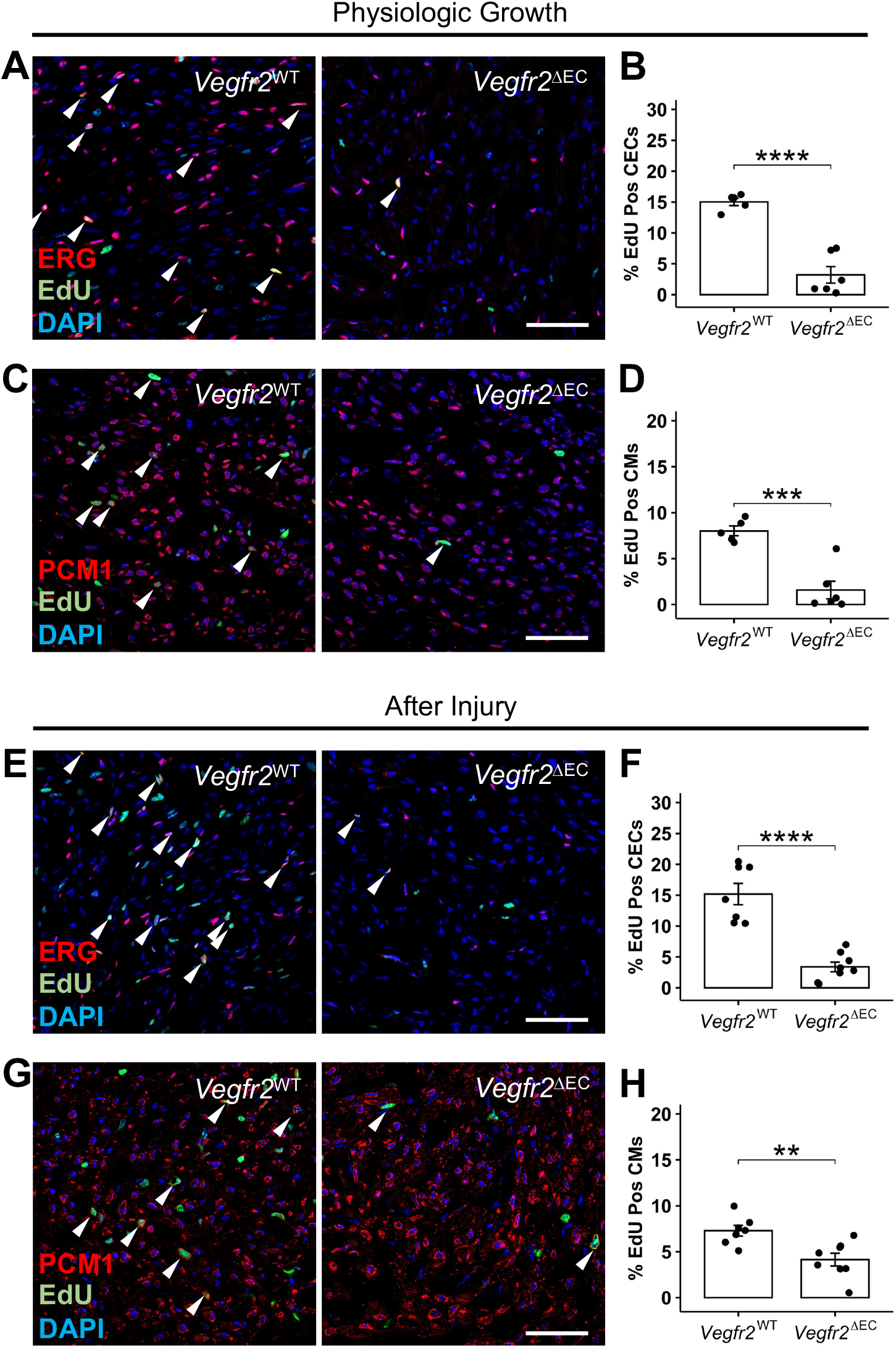
Requirement of *Vegfr2* in CECs for myovascular growth in the neonatal mouse heart. **(A)** Representative images of P4 hearts from *Vegfr2*^WT^ and *Vegfr2*^ΔEC^ mice, immunostained to detect cycling CECs. Arrows point to EdU^+^ Erg^+^ proliferating CECs. (**B)** Quantitation of CEC proliferation in *Vegfr2*^WT^ (n = 5) and *Vegfr2*^ΔEC^ mice (n = 6). ****p = 3.7x10^-5^, two-sided t-test. **(C)** Representative images of P4 hearts from *Vegfr2*^WT^ and *Vegfr2*^ΔEC^ mice, immunostained to detect cycling CMs. Arrows point to EdU^+^PCM1^+^ proliferating CMs. **(D)** Quantitation of CM proliferation in *Vegfr2*^WT^ (n = 5) and *Vegfr2*^DEC^ mice (n = 6). ***p = 0.0004, two-sided t-test. **(E)** Images of P4 hearts from *Vegfr2*^WT^ and *Vegfr2*^ΔEC^ mice after CI at P1. Sections are immunostained to detect cycling CECs. Arrows point to EdU^+^Erg^+^ proliferating CECs. **(F)** Quantitation of CEC proliferation in *Vegfr2*^WT^ (n = 7) and *Vegfr2*^ΔEC^ mice (n = 8). ****p = 2.0x10^-5^, two-sided t-test. **(G)** Representative images of P4 hearts injured at P1 from *Vegfr2*^WT^ and *Vegfr2*^ΔEC^ mice. Sections are immunostained to detect cycling CMs. Arrows point to EdU^+^PCM1^+^ proliferating CMs. **(H)** Quantitation of CM proliferation in *Vegfr2*^WT^ (n = 7) and *Vegfr2*^ΔEC^ mice (n = 8). **p = 0.004, two-sided t-test. Scale bars are 50 μm.

### Inhibition of VEGFA limits CEC and CM proliferation

Our expression data suggest a role for myocardial *Vegfa* in myovascular coupling during growth and after injury (Fig. S1, 3d). To functionally test this concept, we obtained a well-described synthetic antibody, B20-4.1.1 (anti-VEGFA), that binds murine VEGFA and prevents interaction with its receptors (35). Treatment of neonatal mice with anti-VEGFA decreased CEC proliferation by approximately 70% during early neonatal growth and by about 90% after injury indicating a dependency of CEC proliferation on VEGFA (Fig. 5 a,b,e,f). Defects in CEC proliferation following anti-VEGFA treatment were accompanied by decreases in the fraction of EdU^+^ CMs during growth and after injury (Fig. 5 c,d,g,h), further supporting the need for myovascular coupling during growth and regeneration.

**Figure 5:**
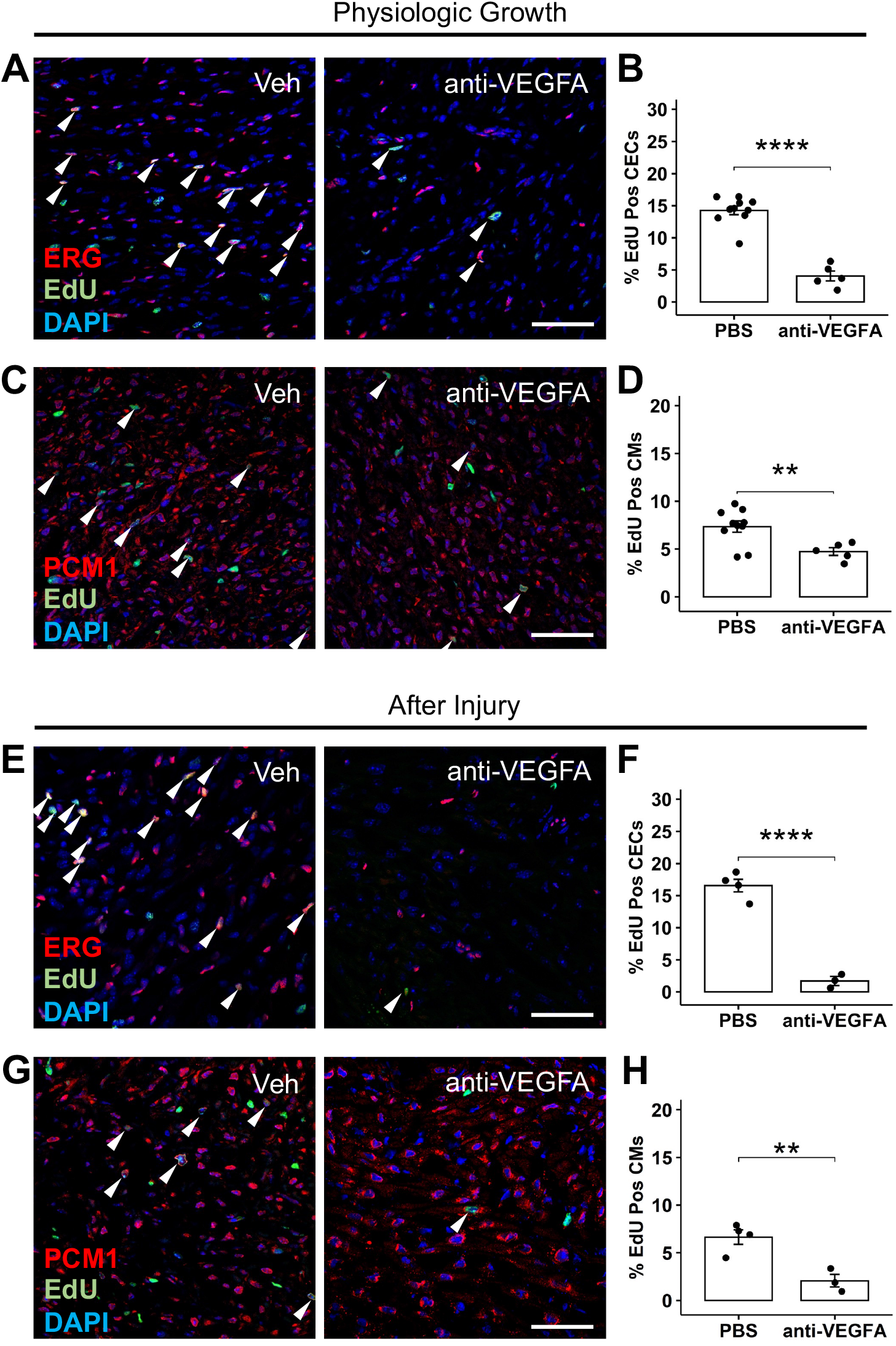
Requirement of VEGFA for myovascular growth in the neonatal mouse heart. **(A)** Representative images of P4 hearts from mice injected with vehicle or anti-VEGFA. Sections are immunostained to detect cycling CECs. Arrows point to EdU^+^Erg^+^ proliferating CECs. **(B)** Quantitation of CEC proliferation at P4 after treatment with vehicle (n = 10) or anti-VEGFA (n = 5). ****p = 1.8x10^-6^, two-sided t-test. **(C)** Representative images of P4 hearts after treatment with vehicle or anti-VEGFA. Sections are immunostained to detect cycling CMs. Arrows point to EdU^+^PCM1^+^ proliferating CMs. **(D)** Quantitation at P4 after treatment with vehicle (n = 10) or anti-VEGFA (n = 5). **p = 0.003, two-sided t-test. **(E)** Images of P4 hearts from mice that underwent CI at P1 and that were injected with vehicle or anti-VEGFA. Arrows point to EdU^+^Erg^+^ proliferating CECs. **(F)** Quantitation of CEC proliferation in *Vegfr2*^WT^ (n = 4) and *Vegfr2*^ΔEC^ mice (n = 3). ****p = 6.9x10^-5^, two-sided t-test. **(G)** Immunostaining for PCM1 and EdU in P4 hearts from mice that underwent CI at P1 and that were treated with vehicle or anti-VEGFA. Arrows point to EdU^+^PCM1^+^ proliferating CMs. **(H)** Quantitation of CM proliferation after treatment with vehicle (n = 4) or anti-VEGFA (n = 3). **p = 0.008, two-sided t-test. Scale bars are 50 μm.

Prior work has described a critical role for tissue hypoxia as a regulator of CM proliferation in both zebrafish and mice (4, 36, 37). Remarkably, hypoxic pre-conditioning of adult mice results in cardiac growth and enhanced regenerative capacity (38). Mechanistically, hypoxia decreases levels of reactive oxygen species, enabling CM cell cycle-reentry (36). As the decreased vascularity in *Vegfr2^ΔEC^* mice and anti-VEGFA treated mice might be expected to result in tissue hypoxia, we examined hypoxia in CMs under these conditions. Pimonidazole is a 2-nitroimidazole used to identify tissue hypoxia based on its formation of stable adducts in the presence of low oxygen tension (39, 40). These adducts can be detected by immunofluorescence with the intensity of staining directly proportional to the level of hypoxia. We administered pimonidazole prior to harvest of hearts from *Vegfr2^ΔEC^* mice, *Vegfr2^WT^* mice, and mice injected with anti-VEGFA or vehicle. We noted an approximately 75% increase in the intensity of pimonidazole uptake by CMs of *Vegfr2^ΔEC^* mice compared to *Vegfr2^WT^* mice and an almost 2-fold increase in mice treated with anti-VEGFA compared to mice treated with vehicle (Supplemental Fig. 3). Together, these results suggest that CECs are a required mediator for CM proliferation in response to tissue hypoxia.

### Exogenous VEGFA enhances the efficiency of innate regenerative responses

Based on the impaired spatial myovascular coupling of CECs and CMs after CI (Fig. 2g-i) and the decrease of *Vegfa* in the border zone after CI (Fig. 3d), we hypothesized that increasing *Vegfa* levels within the border zone might enhance the efficiency of regenerative growth following CI (19). To test our hypothesis, we generated adeno-associated virus to overexpress *Vegfa* (AAV-*Vegfa*) or GFP (AAV-*GFP*). We then cryoinjured mice at P1 and injected AAV-*Vegfa* or AAV-*GFP* into the border zone immediately after injury (Supplemental Fig. 2b-c). Gross examination of hearts injected with a resin to opacify the coronaries hearts at P21 identified increased vascularity at the site of injury of AAV-*Vegfa* hearts compared to AAV-*GFP* hearts (Supplemental Fig. 2c). In addition, we noted that AAV-*Vegfa* hearts had ∼50% less scarring of the left ventricle and better cardiac function after CI (Fig. 6 a-), all suggestive of enhanced regeneration. Importantly, while CEC density was increased in animals treated with AAV-*Vegfa* (Fig. 6 e,f), this was accompanied by a more than 2-fold increase of Ki67^+^ CMs in the border zone compared to hearts treated with AAV-*GFP* (Fig. 6h). Finally, we evaluated sarcomere morphology and *α*-SMA expression, two markers of CM maturity (20, 41). Compared to uninjured hearts and hearts treated with AAV-*GFP*, AAV-*Vegfa* treated hearts were notable for CMs with sarcomeric staining on the periphery of the cell, similar to prior descriptions of sarcomere disassembly (Supplemental Fig. 4a, b) (20). Also consistent with a less mature CM phenotype in AAV-*Vegfa* hearts, CMs were more likely to express the dedifferentiation marker *α*SMA after AAV-*Vegfa* treatment (Supplemental Fig. 4c-e). Together, these data indicate that exogeneous *Vegfa* in the border zone can enhance regenerative responses after neonatal CI by increasing CECs, promoting CM dedifferentiation and proliferation, reducing scarring, and restoring ventricular function.

**Figure 6:**
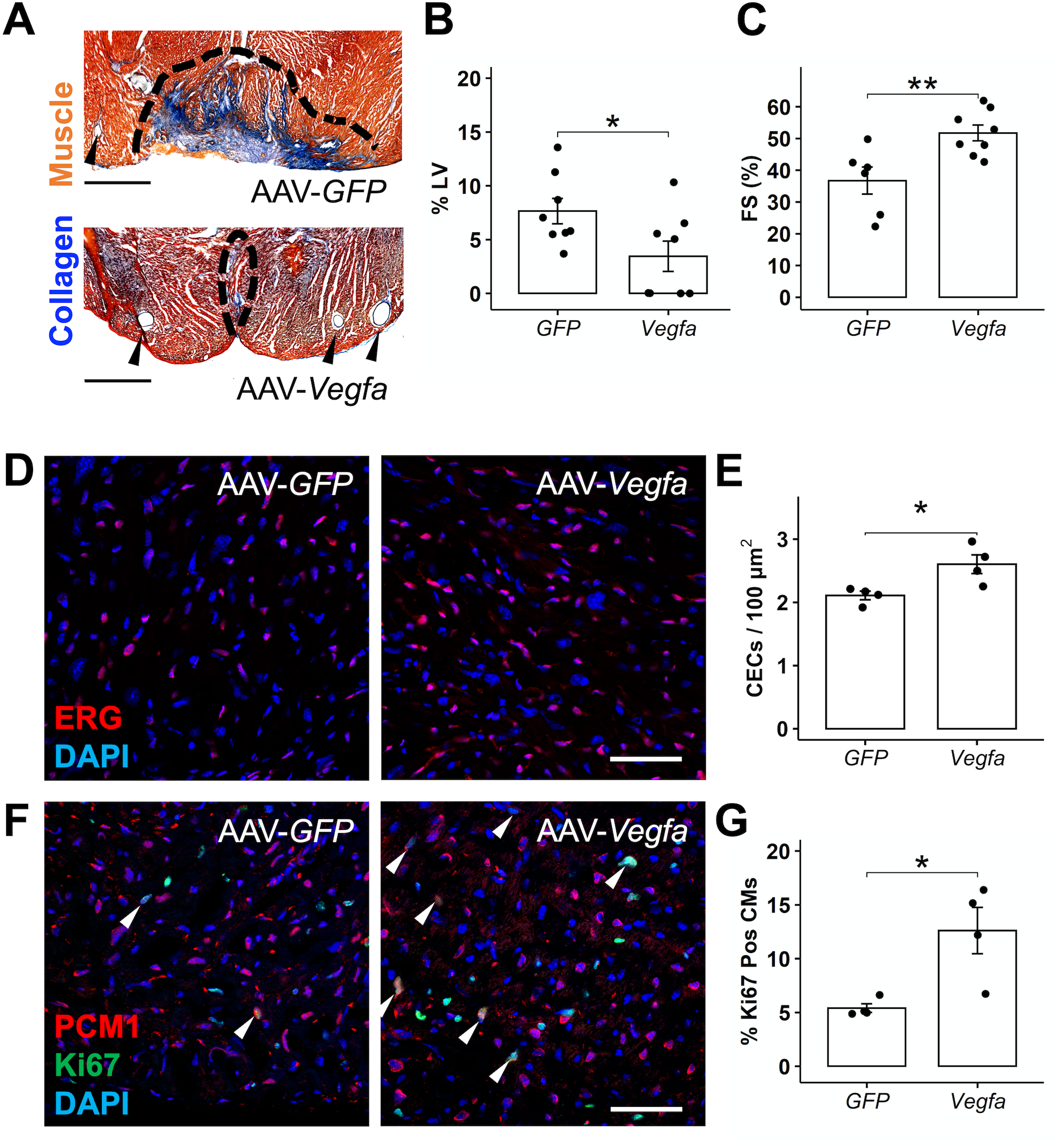
Effect of *Vegfa* overexpression on regeneration after CI. **(A)** AFOG staining of P21 hearts to visualize scar (blue). Arrows indicate vessels. Scale bar is 500 μm. **(B)** Quantitation of scar as a percentage of the left ventricle at P21 in animals injected with AAV-*GFP* (n = 8) or AAV-*Vegfa* (n= 8) at the time of CI at P1. *p = 0.038. **(C)** Fractional shortening at P21 in animals injected with AAV-*GFP* (n=6) or AAV-*Vegfa* (n=8) at the time of CI at P1. **p = 0.008, two-sided t-test. **(D)** Images of the border zone from P10 mice that underwent CI at P1. Sections are immunostained to mark proliferating Erg^+^ Ki67^+^ CECs (arrows). **(E)** Quantitation of CEC density at P14 after injection with AAV-GFP (n=4) or AAV-*Vegfa* (n = 4). *p = 0.035, two-sided t-test. **(F)** Images of the border zone from P14 mice that underwent CI at P1. Sections are immunostained to mark proliferating PCM1^+^Ki67^+^ CMs (arrows). **(G)** Quantitation of CM proliferation at P14 after injection with AAV-*GFP* (n = 4) or AAV-*Vegfa* (n = 4). *p =0.017, two-sided t-test.

### Association of genetically predicted VEGFA levels with myocardial mass in human

Our results after CI in neonatal mice suggest exogenous *Vegfa* could be used therapeutically to promote cardiac growth and regeneration. Indeed, exogenous *Vegfa* has been shown to stimulate cardiac repair after experimental infarction many times in adult mammals, even prompting several human clinical trials in patients with ischemic heart disease (42–47). However, clinical trials have largely failed to improve outcomes, possibly because of inefficient delivery methods that did not sufficiently raise local *Vegfa* levels (48, 49).

Based on its potential as a regenerative factor, we sought to conceptually determine whether VEGFA might regulate cardiac growth in humans using Mendelian randomization (MR). MR is an epidemiologic technique that uses genetic variation to infer causality from observational studies (50–52). Traditional observational studies that associate factors with an outcome are prone to confounding and cannot differentiate causation from reverse causation. However, because innate factors tend to be independent of confounding variables, MR studies are used to point towards causality, and have been used to successfully predict the results of clinical trials (53).

We performed two-sample *cis* Mendelian randomization (MR) analysis with genetically predicted circulating VEGFA concentration as the exposure variable and traits of cardiac structure and function as the outcomes (Fig. 7) (54). Under this framework, genetic variants acting in *cis* that associate with protein abundance (protein quantitative trait loci, pQTL) are used as instrumental variables to estimate the unconfounded causal effects of the protein on the outcomes of interest (Fig. 7a). Genetic variants within the VEGFA gene region were selected from genetic association summary statistics from a genome-wide association study (GWAS) of directly measured circulating VEGFA concentration in over 30,000 individuals (Fig. 7b, Supplemental Table 3) (55). Variant association statistics for six parameters of left ventricular (LV) structure and function were then extracted from a GWAS of cardiac magnetic resonance imaging measures in the UK Biobank (56). A summary of the GWAS is presented in Supplemental Table 4. We applied these instruments, using a two-sample MR model that accounts for partial correlation between instruments, to estimate the effect of genetically predicted circulating VEGFA on LV phenotypes (52). We did not detect a link between genetically predicted VEGFA levels and left ventricular volumes or ejection fraction but found that higher genetically predicted VEGFA levels resulted in increased LV mass (at type I error rate = 0.05 / 6), (Fig. 7c,d). In total, we found an estimated 1.02 gram (95% CI = 0.33-1.70 gram, p = 0.004) increase in LVM per doubling of the circulating VEGFA concentration. Because there is a lack of consensus on how genetic instruments are selected and MR modeling, we performed a sensitivity analysis. Across 120 separate analyses with varying thresholds for instrument selection and four different MR modeling approaches, 119 of these analyses indicate that higher levels of VEGFA are robustly associated with higher LVM (Fig 7e). These results are consistent with the hypothesis that VEGFA promotes cardiac growth in humans and support a possible therapeutic role for VEGFA.

**Figure 7:**
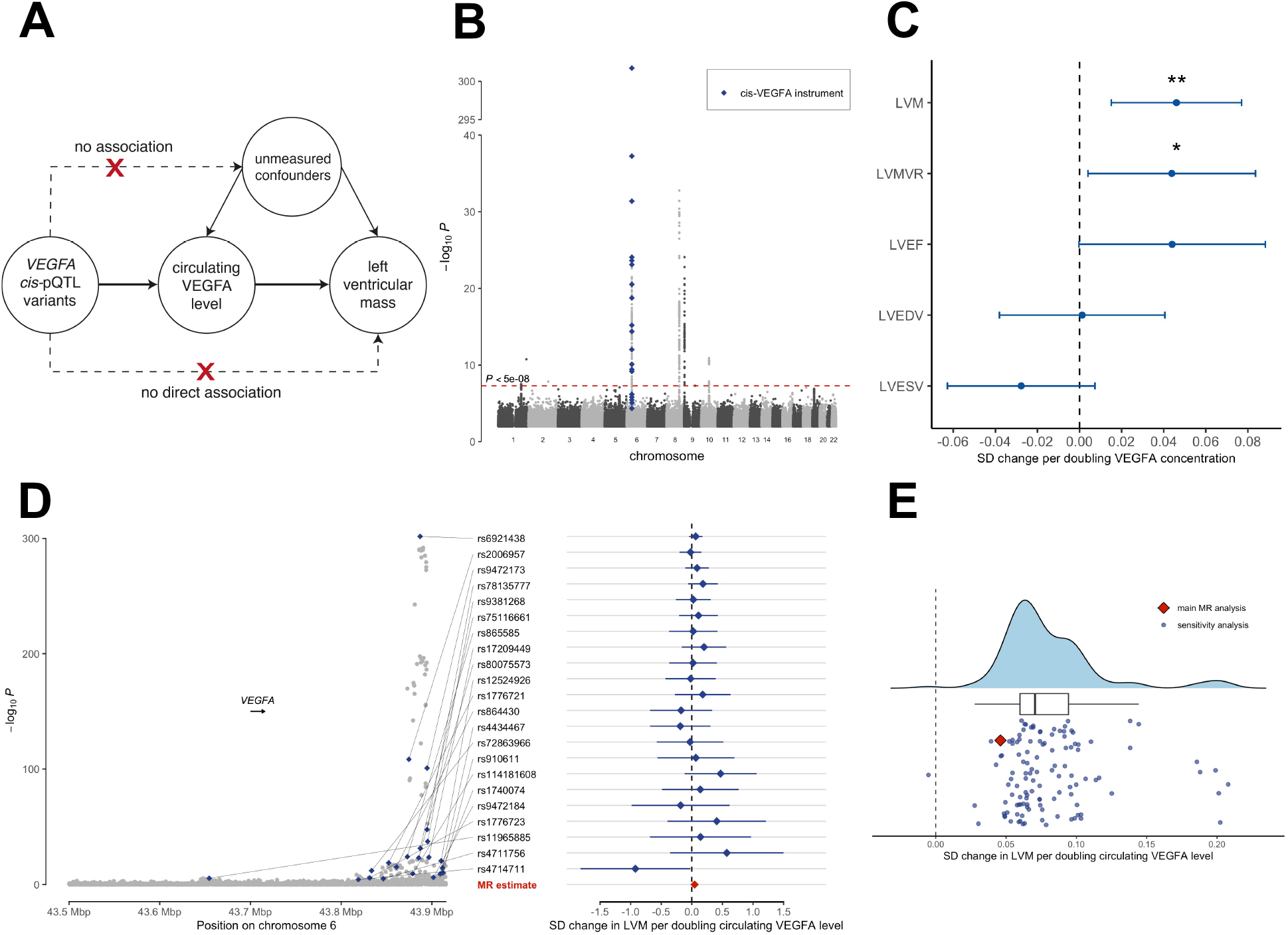
Effect of circulating VEGFA levels on human cardiac structure. **(A)** Illustrative diagram of *cis-*Mendelian randomization (cis-MR) design to estimate the causal association between circulating VEGFA level and left ventricular mass (LVM). **(B)** Manhattan plot showing genome-wide genetic association with circulating VEGFA level. **(C)** *cis*-Mendelian randomization estimate with inverse-variance weighted model of circulating VEGFA level on left ventricular end-diastolic volume (LVEDV), left ventricular end systolic volume (LVESV), left ventricular ejection fraction (LVEF), left ventricular mass to volume ratio (LVMR), and LV mass (LVM). Each data point represents effect estimate in standard deviation (SD) change of the trait per doubling the circulating VEGFA concentration. Error bars indicate 95% confidence intervals (CI) of the estimates. *p < 0.05, **p < 0.01. **(D)** Regional genetic association plot for *cis*-region of the *VEGFA* gene (200 kilobase flanking region from transcript start and end sites) with *cis*-instruments for the main Mendelian randomization analysis highlighted. Estimated causal effects on LVM for each instrument are shown on the right panel with error bars representing 95% confidence interval. **(E)** Distribution of point estimates showing standard deviation (SD) gram change in LVM per doubling circulating VEGFA level as estimated with *cis*-MR with IVW model (red diamond) and sensitivity analyses with permutation of instrument selection methods and *cis*-MR models.

## DISCUSSION

We profiled CEC and CM dynamics to determine that CEC and CM cycling are spatiotemporally coupled in the neonatal mouse heart. Uncoupling CECs from CMs, by deletion of *Vegfr2* from CECs or with an anti-VEGFA antibody, dramatically decreases CM proliferation (Figs 4 and 5). Conversely, improving CEC and CM coupling after CI by increasing local *Vegfa* increases regeneration after CI (Fig. 6). Analogously, myocardial mass of the human heart increases with higher VEGFA levels (Fig. 7). Together, our work demonstrates that the efficiency of cardiac growth is dependent upon myovascular interactions, a finding with strong translational relevance. Approximately 50% of patients with systolic heart failure have flow-limiting coronary artery disease (57). Even in so-called “non-ischemic” cardiomyopathies, cardiac perfusion is impaired due to microvascular disease and capillary rarefaction (58–62). Thus, as methods to promote proliferation of adult mammalian CMs move towards therapeutics, mechanical or molecular revascularization approaches may be a required adjunct for these approaches to be maximally efficacious.

A key concept suggested by our work is that of growth niches, in which cycling CECs establish inductive microenvironments to promote CM proliferation (Fig. 1, 2). We find that cycling CECs are statistically enriched within 7 µm, or 1-2 cell lengths, of cycling CMs during physiologic growth. Because we used pairwise comparisons of CEC and CM distances to deconvolve overlapping niches, our spatial analysis likely underestimates the expanse of the niche surrounding cycling CMs. However, our results are remarkably well-aligned with reported increases of CM proliferation within 3 cell lengths of sprouting angiogenesis in cultured fetal heart sections (63). Niches for CM expansion in the postnatal mammalian heart have been previously identified around hypoxic regions, with hypoxia even being able to induce CM proliferation in the adult mouse heart (4, 38). Mechanistically, hypoxia exerts its effect on CM proliferation cell autonomously, by modulating reactive oxygen species and the DNA damage response (36). Our work suggests another element to the phenomenon of hypoxia-induced CM proliferation. We were able to induce hypoxia in the neonatal heart through two different approaches, genetic deletion of *Vegfr2* from CECs and with an anti-VEGFA antibody (Supplemental Fig. 3). However, even though hypoxia would be predicted to increase CM proliferation, we observed decreased CM cycling in these models. Thus, hypoxia mediated CM expansion is likely to involve both CM-specific effects but to also depend on CECs within the hypoxic niche. While hypoxia may be one approach to stimulate heart regeneration, therapeutic hypoxia for patients with heart failure may be challenging. A better understanding of the signaling milieu of the hypoxic niche could lead to alternate approaches for stimulating hypoxia-mediated regeneration.

Because depletion of CECs, either through deletion of *Vegfr2* from CECs or by administration of anti-VEGFA, leads to defects in CM proliferation, our work demonstrates that CECs promote CM cycling (Fig. 5, 6). Mechanistically, CEC expansion may simply lead to new conduits for growth factors or other cell types that promote cardiac growth. However, multiple lines of evidence also point to a direct role for CECs to influence CM proliferation through angiocrines. Indeed, recent work in zebrafish and neonatal mice support this concept. In zebrafish, inhibition of Notch in CECs results in reduced secretion of Wnt inhibitors that influence CM proliferation (64). More directly, deletion of *Igf2* from CECs in developing mice abrogates CM proliferation after neonatal cardiac injury (65). Interestingly, deletion of *Igf2* from CECs does not affect physiologic CM cycling, raising the intriguing possibility that different angiocrines modulate myocardial expansion during physiologic growth and after injury. Finally, different CEC subsets may contribute different sets of angiocrines, as recent work suggests that *Reln* from lymphatic endothelial cells has direct effects on CM proliferation (13). Future work to identify the complement of angiocrines that instruct CM proliferation in different developmental and injury contexts is needed.

Our work also suggests that restoring myovascular coupling can enhance regenerative capacity. Unlike physiologic growth, CEC cycling is not enriched around cycling CMs in the border zone after CI, a model of incomplete regeneration. Functionally increasing CEC density through overexpression of the master angiokine *Vegfa* enhances CM cycling and reduces scarring after CI, both signs of enhanced regeneration (Fig. 6). Taken together with our prior work showing that *vegfaa* overexpression can induce ectopic heart regeneration in zebrafish, exogenous *Vegfa* may be one approach to augment regeneration. Indeed, our MR study indicates that VEGFA levels are causally linked to human myocardial mass and predict that more VEGFA can increase human cardiac growth (Figure 7). Several clinical trials of *Vegfa* are now underway to revisit whether exogenous VEGFA can be used therapeutically to treat human cardiovascular disease (NCT03409627, NCT03370887, and NCT04125732). Compared to prior clinical trials, these studies utilize newer delivery methods such as modified RNAs, newer generation viral vectors, and more efficient plasmid delivery systems. Based partly on our findings, we would expect that markers of increased CECs, such as improvements in myocardial perfusion, will predict VEGFA-based treatment effects on cardiac mass and function.

We acknowledge several limitations to our work. First, although we focus on the coupling of CECs and CMs during growth and regeneration, additional cell types also regulate cardiac growth and are likely to be present within niches. Defining niche constituents may be critical to efficiently instruct cardiac growth. Second, while we demonstrate a critical role of VEGFA-VEGFR2 signaling to coordinate CEC and CM expansion, VEGFR2 broadly marks the cardiac microvasculature. Future work to refine which microvascular cells instruct CM cycling could enable more targeted regenerative approaches via cell transplantation or specific signaling cues. Finally, while our MR studies point to a therapeutic benefit to using VEGFA to stimulate cardiac growth, our MR studies model a lifetime of exposure to VEGFA and do not have the resolution to confirm that increased myocardial mass is the result of CM expansion. Additional work to determine whether VEGFA promotes cardiac growth across developmental stages and to directly link VEGFA to human CM proliferation in vivo are needed to establish VEGFA as a bona fide regenerative factor in humans.

In summary, here, we present work showing that coordinated growth of CECs and CMs guides postnatal cardiac growth and regeneration. We speculate that a better understanding of the signaling milieu within a myovascular niche can inform approaches for heart regeneration.

## METHODS

### Mice and Neonatal Cryoinjuries

CD-1 mice (Charles River Labs, Morrisville, NC) were used for profiling myovascular kinetics during growth and regeneration. CD-1 mice were also used for anti-VEGFA antibody and AAV experiments. *Cdh5-CreER*^T2^ and *Vegfr2^flox/flox^* strains have been previously described (33, 34). *Cdh5-CreER*^T2^; *Vegfr2^flox/flox^* and *Vegfr2^flox/flox^* mice were treated with 100 mg of tamoxifen intraperitoneally at P0 and P1. Complete recombination was verified by immunostaining for VEGFR2 (Supplementary Fig. 2). For EdU incorporation studies, 0.5 mg EdU was given intraperitoneally 4 hours prior to tissue harvest. Cryoinjuries were performed on P1 pups as previously described (25, 28).

The number of animals used in these studies is specified in the figure legends. For mouse experiments, we included animals 1) confirmed to have *Vegfr2* deletion by immunostaining for VEGFR2; 2) animals with weight loss after anti-VEGFA treatment; or 3) animals confirmed to have *Vegfa* overexpression by in situ hybridization after injection with AAV-*Vegfa*. For assays without computational scoring, readers were blinded to experimental groups. All protocols were approved by the Institutional Animal Care and Use Committee at Duke University.

### Histology and Immunostaining

At the time of tissue harvest, hearts were perfused with KCl and then 4% PFA. Hearts were then immersed in 30% sucrose overnight and embedded in Tissue Freezing Media for cryosectioning.

Cryosections were blocked with PBST (PBS with 0.1% Tween-20) containing 10% newborn calf serum and 1% DMSO and incubated overnight with primary antibodies at 4°C. Cryosections were then washed with PBST and incubated with secondary antibodies and DAPI (100 ng/ml). For EdU detection, cryosections were incubated in EdU staining solution (100 mM Tris-HCl, 1 mM CuSO_4_, 10 mM Azide, and 50 mM ascorbic acid in PBS) for 10 minutes. Primary antibodies used for this study included: anti-PCM1 (Sigma, HPA023370, 1:100); anti-PCM1 (Santa Cruz, sc-398365, 1:100); anti-Erg (Abcam, ab92513, 1:25); anti-CD31 (BDBioscience, 553370, 1:100), anti-VEGFR2 (BDBioscience, 555307, 1:100), anti-Ki67 (ThermoFisher, 4-5698-82, 1:100), anti-Actinin (Cell Signaling, 6487P, 1:100), anti-Tnnt (DSHB CT3, 1:25) *α*SMA (Abcam, ab5694, 1:200). Secondary antibodies and azides were conjugated to Alexa-488, Alexa-594, or Alexa-633 (Invitrogen). To label CM nuclei, cryosections underwent heat induced epitope retrieval with citrate buffer prior to immunostaining. Stained slides were imaged on a Zeiss AxioImager M1 epifluorescent microscope, a Zeiss CSU-X1 spinning disk confocal microscope, or a Zeiss LSM 510 confocal microscope. For physiologic growth experiments, 9 -16 non-overlapping images (40X) of the left ventricle were obtained for each section. For CI experiments, the border zone was defined as the entire region within 1-2 40X fields of view along the entirety of the injury plane. For each heart, the 3 largest sections were imaged. **Quantification of CM and CEC Proliferation**

We developed several customized image segmentation pipelines using CellProfiler and Ilastik for automated scoring EdU^+^ CECs and CMs (66, 67). For CEC quantification, grayscale images for each channel were processed with rolling-ball background subtraction and used as input into a CellProfiler routine that 1) identified Erg^+^ nuclei based on Erg staining and DAPI intensity; 2) identified EdU^+^ nuclei based on EdU staining and DAPI staining; and 3) identified EdU^+^ Erg^+^ nuclei based on the presence of an EdU object within an Erg object.

For CMs, grayscale images from individual channels were obtained using a Zeiss CSU-X1 spinning disk confocal microscope. Images were pre-processed to generate 32-bit grayscale images and to create a set of images with merged PCM1 and DAPI channels. The DAPI image and the merged PCM1-DAPI images were then input into machine learning routines to generate probability maps for DAPI^+^ nuclei and PCM1^+^/DAPI^+^ nuclei, using Ilastik. Machine learning algorithms were trained using images from our physiologic growth experiments. Probability maps and grayscale images were used as input for a CellProfiler pipeline that 1) identified nuclei based on a DAPI probability map; 2) filtered nuclei for CMs based on the mean PCM1^+^/DAPI^+^ pixel probability; 3) identified EdU^+^ nuclei based on EdU staining and DAPI staining; and 4) identified EdU+ CM nuclei based on the presence of an EdU object within a CM nucleus. CellProfiler output data was tabulated using the dplyr package in R (68).

### Proximity mapping

For proximity mapping, z-stack images of P4 hearts were obtained using a Zeiss LSM 510 confocal microscope. CM and CEC nuclei were segmented using the Spots tool along with manual refinement, assigned coordinates, and categorized as EdU^+^ or EdU^-^ using Imaris (Bitplane AG, Zurich, Switzerland). Coordinate files were then used to determine the pairwise distance of each CM and each CEC nucleus. Distances were used to compute the density of EdU^+^ CECs as a function of distance from each PCM1^+^ nucleus using the dplyr and plotly packages in R (68, 69).

### In situ hybridization

RNAscope Probe - Mm-Vegfa-O2 (Lot 16197A), with predicted reactivity against all known murine *Vegfa* variants, was hybridized against cryosections using the manufacturer’s protocol (Advanced Cell Diagnostics, Hayward, CA). Images of sections were quantified using a CellProfiler routine adapted from Erben et al (32). Briefly, the DAPI channel was used to identify nuclei objects and the number of *Vegfa* spots adjacent to each nucleus was counted.

### Single Cell RNA Sequencing and Analysis

The left ventricles of cryoinjured hearts were collected at P7. The apical third of the LV that contained the injured area was then dissociated into single cells. Single cells were then captured into droplets using microfluidics followed by preparation of single cell cDNA libraries as previously described (70). Samples were sequenced on a single lane of an Illumina HiSeq. Sequencing reads were mapped to Ensembl release NCBIM37.67. Aligned reads were binned and collapsed onto cellular barcodes using the Drop-seq pipeline v1.13.3 (http://mccarrolllab.com/dropseq), resulting in a raw digital expression matrix containing the count of unique UMIs for each gene in each cell (71).

Expression analysis was performed by the Duke Bioinformatics Shared Resource, using the Seurat package (v2.2.0) in R/Bioconductor (72). Cells with gene counts over 4,000 or less than 200 were removed. Genes expressed in less than three cells were also removed. To avoid over-filtering CMs, we did not filter cells based on high expression of mitochondrial genes. Gene expression, for 1721 cells across 12754 genes, was normalized by scaling by the total number of transcripts, multiplying by 10,000, and log transformation. Unwanted sources of variation were adjusted for by regression on the number of detected UMIs using the *ScaleData* function. We used the JackStraw method to determine the number of statistically significant principal components (PCs). The *FindClusters* function was used to identify cellular clusters, using the 20 significant PCs and a 0.6 resolution. We used the t-SNE method to visualize cells across 20 PCs in two dimensions. To identify cell types, we used the *FindAllMarkers* function. Parameters were set to test all genes, expressed in at least 10% of the cells in each cluster, for differential expression.

### Anti-VEGFA Antibody Treatment

Anti-VEGFA antibody (B20-4.1.1) was a kind gift from Genentech (San Francisco, CA). Neonatal mice were injected with 5 mg/g of anti-VEGFA or vehicle at P0 and P4. Adequate injection of anti-VEGFA was confirmed by significant weight loss compared to control animals at the time of tissue harvest.

### Determination of Hypoxia

Mice were given 60 mg/g pimonidazole (Hypoxyprobe, HP3-100kit) intraperitoneally 1.5 hours prior to tissue harvest. Tissues sections were stained with an anti-pimonidazole antibody (Hypoxyprobe, HP3-100kit, 1:100) and an antibody against a-actinin. For each heart, 9 images of the left ventricle for 3 separate cardiac sections were obtained using an AxioImager M1 microscope. Images were quantified using a CellProfiler routine to 1) create an image mask based on the a-actinin stain and 2) determine the mean intensity of anti-pimonidazole stain.

### AAV Generation and Treatment

AAV encoding *Vegfa-164* under the control of a CMV promoter was generated by PCR amplifying the *Vegf164* isoform of *Vegfa* from a mouse endothelial cDNA library and cloning this fragment into the pTR2-eGFP with replacement of *EGFP* (73). AAV, serotype 9, was generated through the Duke Viral Vector core. AAV virus encoding *GFP* under the control of a CMV promoter was obtained from the Duke Viral Vector Core. For intracardiac injections, 12 μl of AAV at 1x10^9^ vg/ml was injected circumferentially around the injured area immediately after CI (Supplemental Fig. 2b).

### Coronary vessel labeling

The coronary vasculature was visualized by injection of a low viscosity polyurethane resin (PU4ii, VasQTec) into the apex of the left ventricle (74). Whole mount images were obtained using a stereoscope.

### Scar Assessment

P21 hearts were serially sectioned from base to apex and stained with acid fuchsin orange G, as previously described (75). Stained slides were scanned with a Leica Aperio Digital Pathology Slide Scanner. Scar area as a percentage of the left ventricle was scored by a blinded reader. The average scar percentage across the 5 largest sections was determined for each heart.

### Echocardiography

Fractional shortening was determined by echocardiography using the Duke Cardiovascular Physiology Core. Images were analyzed by three blinded readers and an averaged reading is reported.

### Mendelian Randomization to Determine Effects of Circulating VEGFA Levels on Human Cardiac Structure

Two-sample MR analysis was performed using summary statistics from GWAS of circulating protein level measured with Olink proximity-extension assay in 30,931 individuals of European ancestries and GWAS of cardiac MRI – derived left ventricular phenotypes in 16,923 UK Biobank participants (Supplementary Table 4) (55, 56). To represent effect in the original unit of measurements, the standardized estimates from both GWAS were back-transformed by multiplying the MR effect estimate with the estimated standard deviation of the traits. MR instruments for VEGFA were selected from variants that 1) are available in both GWAS, 2) located within the 200 kilobases flanking region from the genomic coordinate of *VEGFA* gene; 3) have a MAF > 0.01; 4) have a *P*-value for association with circulating VEGFA level < 1x10^-4^; and 5) are LD-clumped to an *r^2^* threshold of 0.4 using *plink* with --clump option (76). Mendelian randomization analysis was performed using an inverse-variance weighted model accounting for correlation between instruments, implemented in the *MendelianRandomization* R package (77, 78). Correlation between instruments were estimated from a random sample of 10,000 UK Biobank participants of European ancestries (79).

To test the robustness of the cis-MR analysis, a series of sensitivity analyses were performed by varying parameters for instrument selection and MR models. For instrument selection, LD clumping was performed using five different *r^2^* thresholds (0.05, 0.1, 0.2, 0.4, and 0.6) and six different *P*-value thresholds for association with circulating VEGFA level (no threshold), 1e^-2^, 1e^-3^, 1e^-4^, 1e^-5^, and 5e^-8^). Four different MR models were also tested for each instrument set (inverse-variance weighted, MR-Egger, Principal component MR with 90% variance explained, and Principal component MR with 99% variance explained) (80, 81). To account for correlation between instruments, we implemented the method proposed by Burgess *et al.* using an instrument correlation model derived from genotype data of a random subset of 10,000 UK Biobank participants of European ancestry (77).

### Statistics

All means are presented as mean ± SEM and proportions as proportion ± 95% confidence interval (95% CI). Statistical analysis between two groups was performed using a two-tailed unpaired t-test test. Pairwise comparisons between multiple groups were performed using a two-tailed unpaired t-test test with correction for multiple testing using Holm’s method. A p < 0.05 was set as an *a priori* threshold for significance. Statistical analysis and plots were generated in R using the dplyr, ggpubr, and ggplot2 packages (68, 82, 83). For each experiment, individual data points are presented in the plot and the sample size is specified in the figure legend.

### Study approval

All animal protocols were approved by the Institutional Animal Care and Use Committee at Duke University.

## Acknowledgments

We thank Ken Poss, Nenad Bursac, Doug Marchuk, and Howard Rockman for comments and discussion. We would like to thank Dr. Helene Fradin Kirshner from the Duke Bioinformatics Shared Resource for assistance with the analysis of scRNA-seq data.

## Author contributions

Conceptualization: RK Methodology: RK, RTL, PRT, SHS

Investigation: PD, AK, AH, KB, AP, MCT, YK, TM, CK, RTL, RK

Visualization: PD, AK, AH, RTL, RK

Funding acquisition: RK, RTL

Project administration: PD, RTL, RK

Supervision: RK, RTL

Writing – original draft: PD, RK

Writing – review & editing: PD, AH, SHS, CDK, RTL, RK.

## Sources of Funding

National Institutes of Health grant R03 HL144812 (RK)

Duke University Strong Start Physician Scientist Award (RK),

Mandel Foundation Seed Grant (RK)

Walker P. Inman Endowment (RK)

UK Research and Innovation Rutherford Fellowship MR/S003754/1 hosted by Health Data Research UK (RTL)

BigData@Heart Consortium funded by the Innovative Medicines Initiative-2 Joint Undertaking grant 116074 (RTL)

National Institute for Health Research University College London Hospitals Biomedical Research Centre (RTL)

---

**Figure S1.**
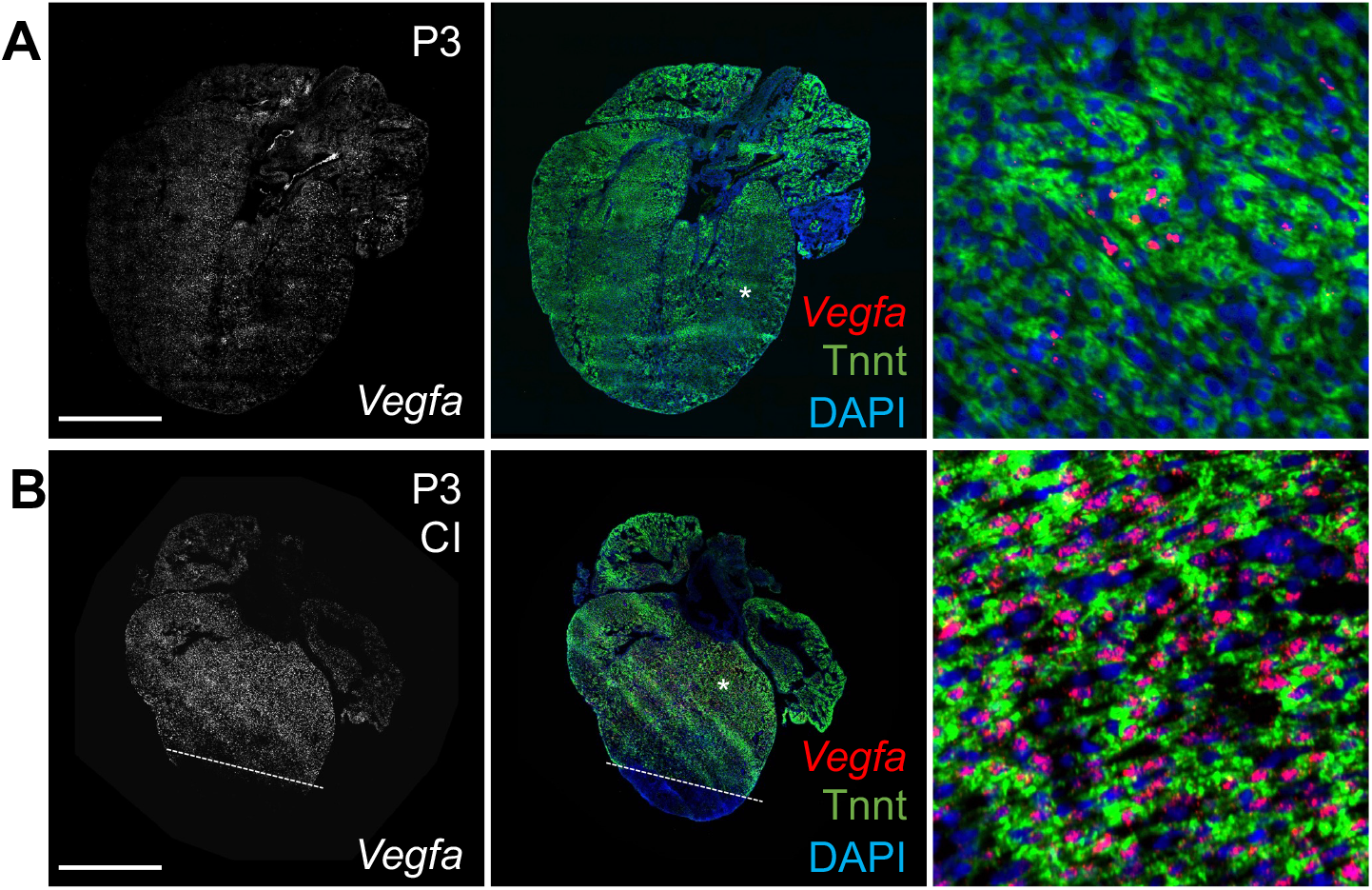
*Vegfa* expression in the neonatal mouse heart. **(A-B)** Tilescan images of an uninjured P3 heart (A) and a P3 heart that underwent CI at P1 (B). Sections are immunostained for Tnnt (green) and hybridized with an in situ probe for *Vegfa* (red). Lines correspond to the approximate injury plane. The asterisks correspond to the magnified images. Scale bar is 500 μm.

**Figure S2.**
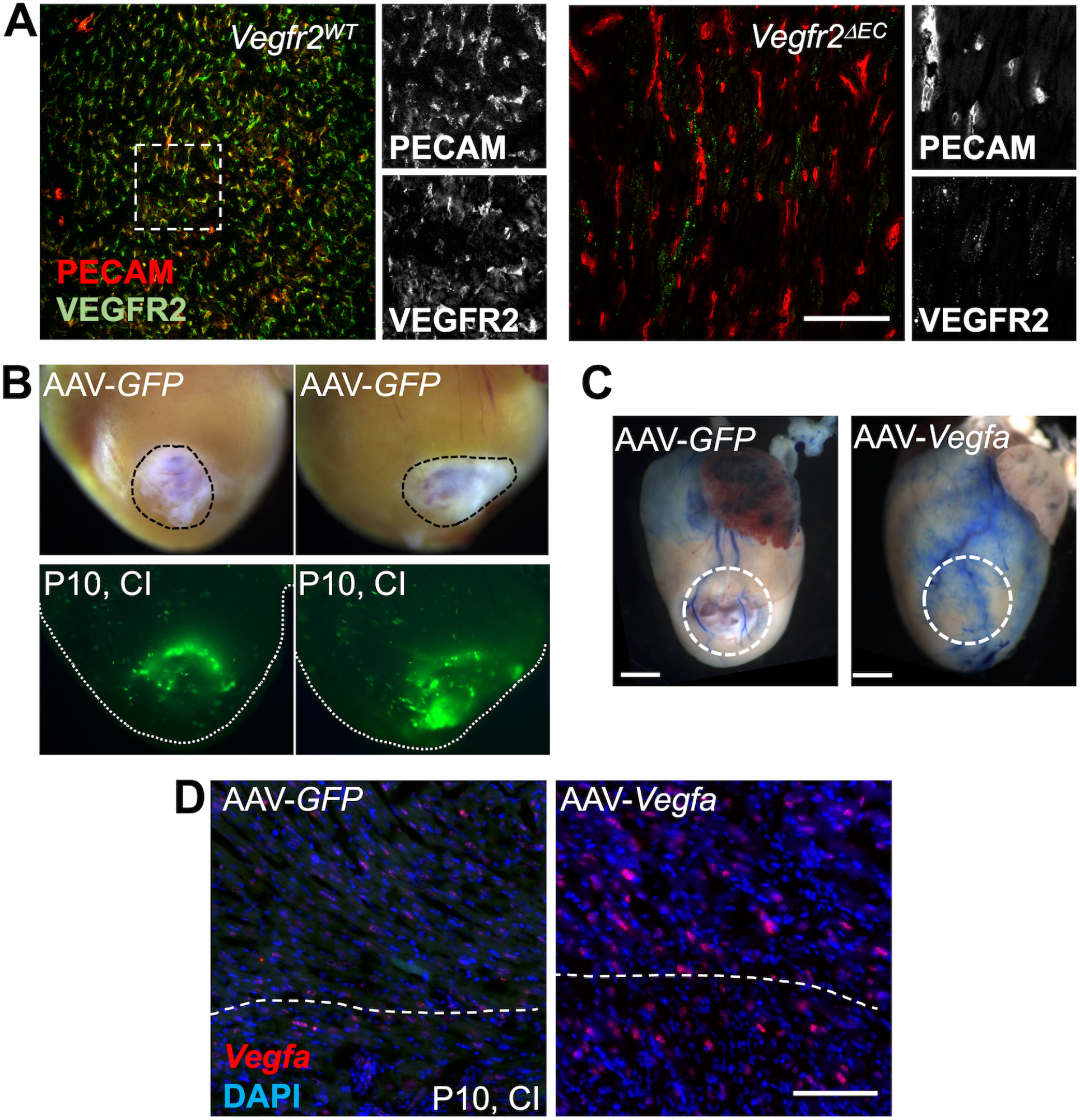
Approaches to manipulate VEGFA/VEGFR2 signaling. **(A)** Images of *Vegfr2*^WT^ and *Vegfr2*^ΔEC^ mice at P3 after treatment with tamoxifen at P0. Sections are immunostained for PECAM and VEGFR2. The dashed box corresponds to magnified images, shown as individual channels. Scale bar is 100 μm. **(B)** Whole mount images of representative P10 hearts injected with AAV-*GFP* at the time of CI at P1. (Top) Brightfield images of the ventricles with the black dashed circle indicating the area of injury. (Bottom) Corresponding GFP images to show concentration of GFP expression around the area of injury. **(C)** Whole mount images of representative P21 hearts, injected with either AAV-*GFP* or AAV-*Vegfa* at the time of CI at P1. Blue resin opacifies the coronary vasculature. Circled area indicates the approximated area of injury. Scale bar is 1 μm. **(D)** Single molecule in situ hybridization for *Vegfa* in the border zone of P10 hearts that were injected with either AAV-*GFP* or AAV-*Vegfa* after CI at P1. Dashed line indicates the approximate plane of injury, with the region above the line corresponding to the border zone. Red dots indicate *Vegfa* expression. Scale bar is 50 μm.

**Figure S3:**
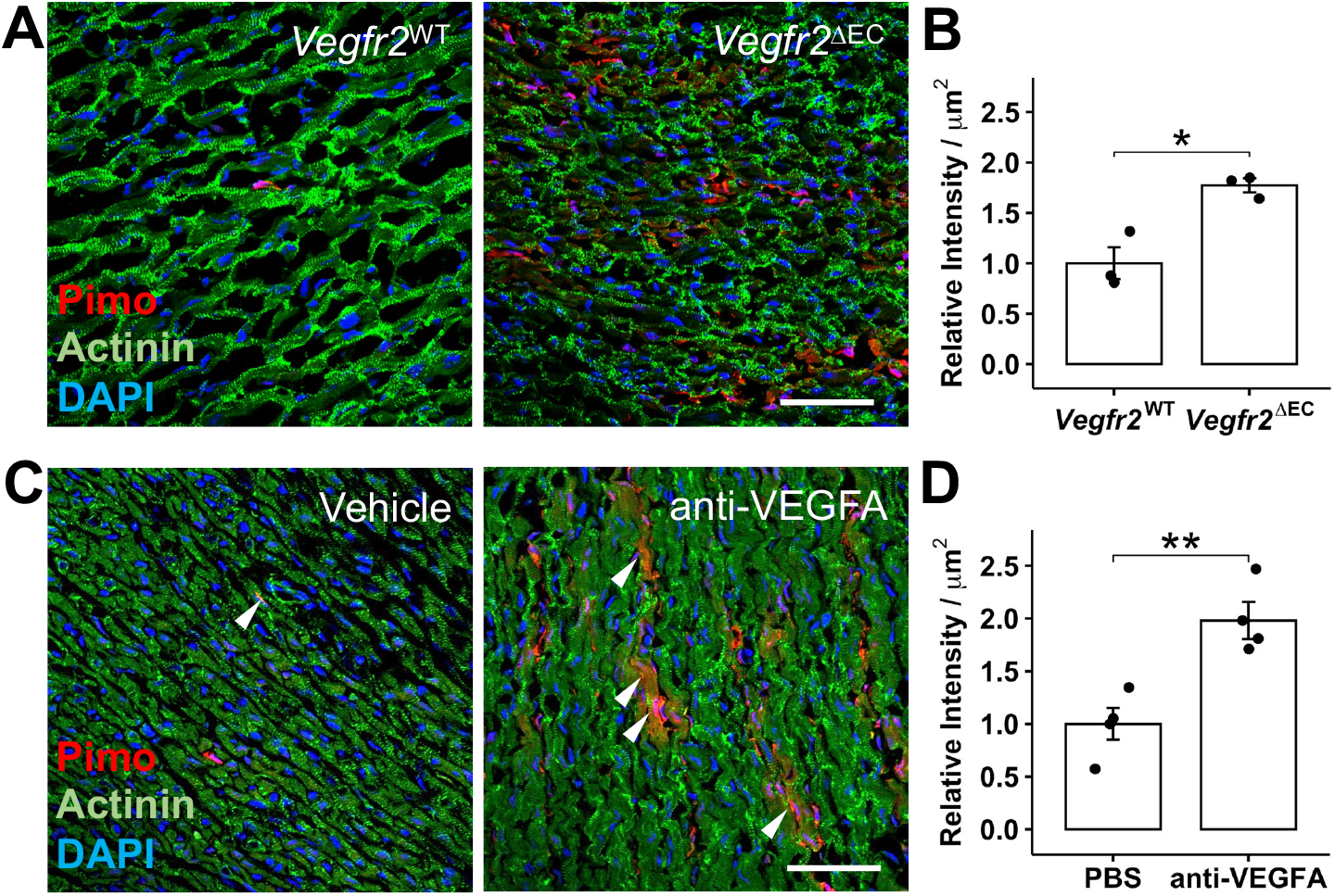
Myocardial hypoxia with loss of *Vegfr2* in CECs or after administration of anti-VEGFA. **(A)** Images of sections from P4 *Vegfr2*^WT^ and *Vegfr2*^ΔEC^ hearts immunostained for actinin (green) and pimonidazole (red) incorporation. **(B)** Quantitation of pimonidazole intensity per μm of tissue stained with actinin in *Vegfr2*^WT^ (n = 3) and *Vegfr2*^ΔEC^ (n = 3) hearts. *p = 0.011, two-sided t-test. **(C)** Images of sections from P4 mice treated with vehicle or anti-VEGFA. Sections are immunostained for actinin (green) and pimonidazole (red) incorporation. **(D)** Quantitation of pimonidazole intensity per μm of tissue stained with actinin in mice treated with vehicle (n = 4) or anti-VEGFA (n=4). **p = 0.005, two-sided t-test. Scale bars are 50 μm.

**Figure S4:**
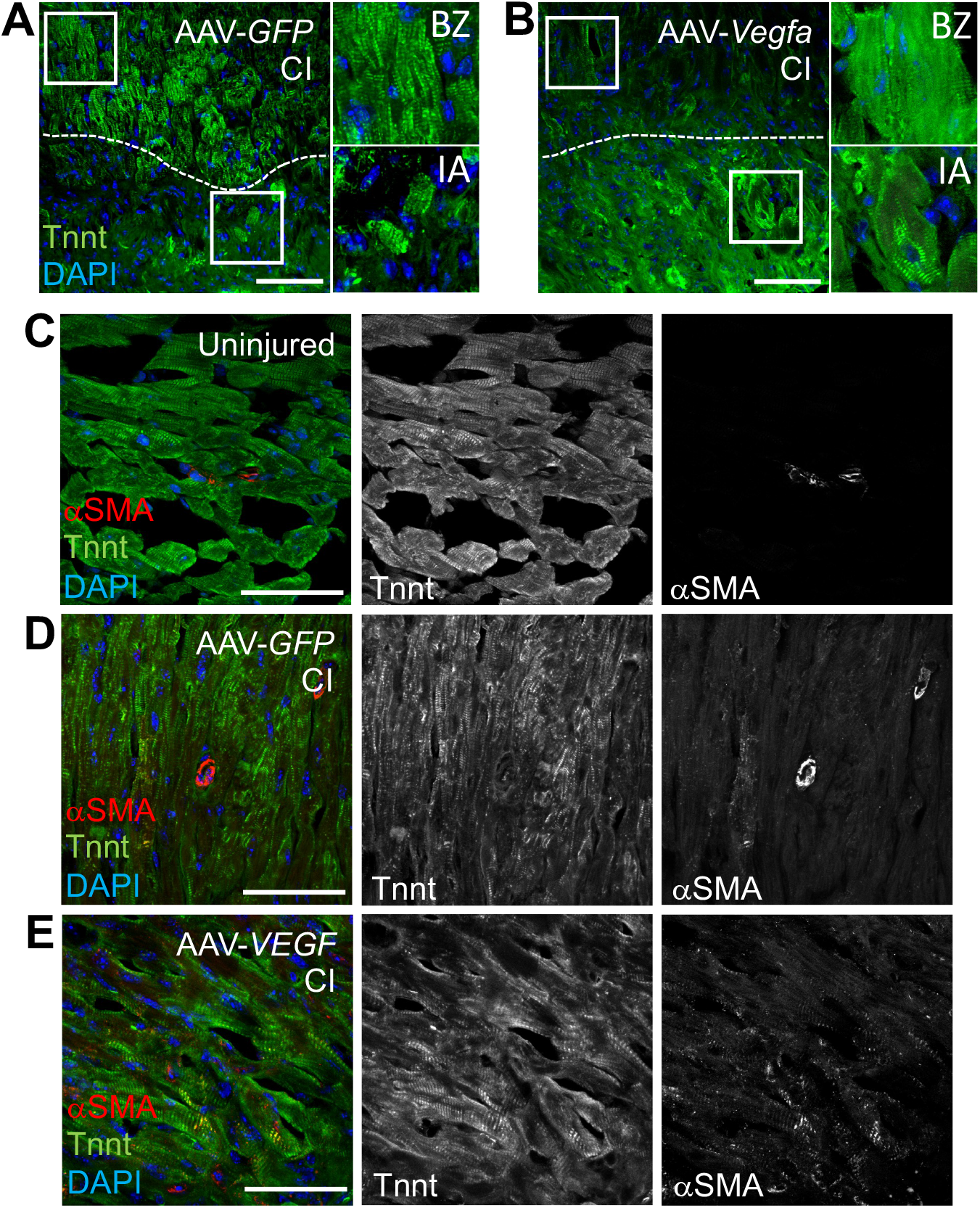
Effect of *Vegfa* overexpression on sarcomere morphology and *α*SMA expression in CMs after CI. **(A-B)** Images from the site of injury of P21 mice that were treated with AAV-*GFP* or AAV-*Vegfa* at the time of CI at P1. Dashed line indicates the approximate plane of injury. The region above the line is the border zone (BZ) and the region below the line is the injured area (IA). Boxed regions correspond to the adjacent magnified panels. Sections are immunostained for the sarcomeric protein Tnnt (green). **(C-E)** Images of the border zone from P21 mice that underwent injection of AAV-*GFP* **(D)** or AAV-*Vegfa* **(E)** at time of CI at P1 or physiological control **(C)**. Sections are immunostained to mark CMs (Tnnt, green) and alpha-smooth muscle actin (*α*SMA, red). Scale bars are 50 μm.

